# Serological investigation of influenza D virus in cats and dogs in Europe

**DOI:** 10.64898/2026.01.03.697426

**Authors:** Claudia Maria Trombetta, Aurora Fiori, Alessandro Falsini, Francesco Pellegrini, Sophie Le Poder, Amit Eichenbaum, Venetia Cardona, Justine Oliva, Gilles Meyer, Nataliia Muzyka, Denys Muzyka, Emanuele Montomoli, Barbara di Martino, Mariette F. Ducatez, Gianvito Lanave, Vito Martella, Michele Camero

## Abstract

A multicentric European study investigated the seroprevalence of influenza D virus (IDV) in domestic cats and dogs. Serum samples from Italy, France and Ukraine (2015-2018, 2020, 2023-2024) showed no IDV positivity in Italy or in France. In Ukraine, 2.46% of dogs and 0.85% of cats tested positive in 2024.

---

Influenza D virus (IDV) has cattle as its primary reservoir but can occasionally spill over into other species (1, 2). Recent serological findings in dogs (Italy) and cats (China) suggest a broader host range than previously recognized (3, 4).

In this study, we investigated the IDV seroprevalence in feline and canine samples in a multicentric European study (Italy, France and Ukraine).

Serum samples from domestic cats were collected in 2023 (n=76) and 2024 (n=56) in the Apulia region, Italy by veterinary offices either for presurgical evaluation or for routine analysis. In France, serum samples (n=114 dogs and n=47 cats), obtained from the “companion animal” clinic at the École Vétérinaire de Maisons-Alfort (ENVA, Paris, Ile-de-France region), were collected between 2015 and 2018 during animal hospitalization. Dog nasal swabs (n=41) and lung tissues (n=24) either originated from a shelter or from the ENVA clinics from animals with respiratory clinical signs.

In Ukraine, serum samples from domestic cats (n=118) and dogs (n=122) were collected in 2020, 2023 and 2024 in veterinary clinics from Donetsk, Zaporizhzhia, Khmelnytska, Odesa, Kyiv, Lviv, Kharkiv, and Dnipropetrovsk Oblasts (table 1 and Supp figure 1).

**Table 1.**
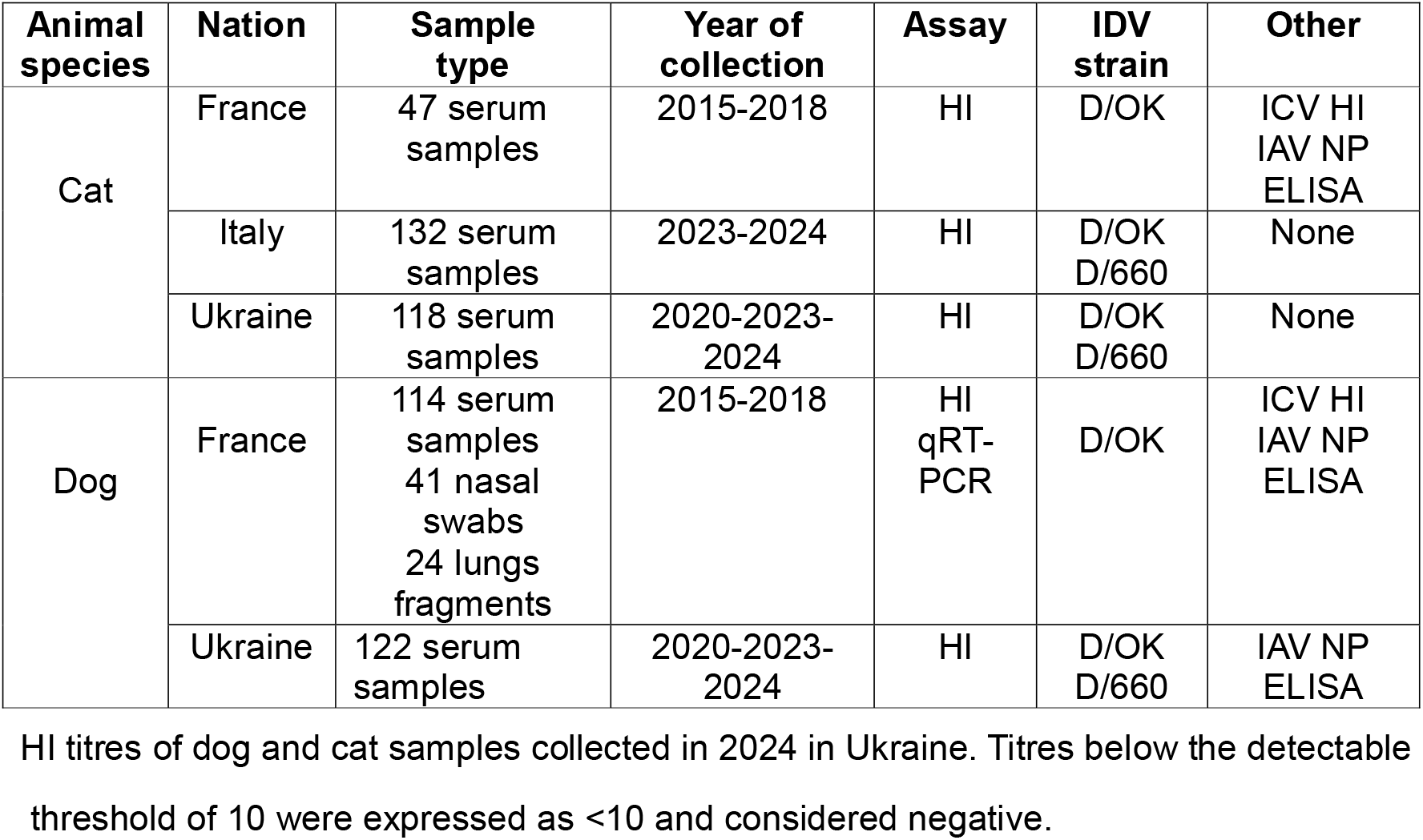
Overview of the cat and dog samples included in the study, along with information on the country, year of collection, testing assays, IDVs strains and other additional information.

| Animal species | Nation | Sample type | Year of collection | Assay | IDV strain | Other |
| --- | --- | --- | --- | --- | --- | --- |
| Cat | France | 47 serum samples | 2015-2018 | HI | D/OK | ICV HI<br>IAV NP<br>ELISA |
|  | Italy | 132 serum samples | 2023-2024 | HI | D/OK<br>D/660 | None |
|  | Ukraine | 118 serum samples | 2020-2023-2024 | HI | D/OK<br>D/660 | None |
| Dog | France | 114 serum samples<br>41 nasal swabs<br>24 lungs fragments | 2015-2018 | HI<br>qRT-PCR | D/OK | ICV HI<br>IAV NP<br>ELISA |
|  | Ukraine | 122 serum samples | 2020-2023-2024 | HI | D/OK<br>D/660 | IAV NP<br>ELISA |

All samples were tested in duplicate by hemagglutination inhibition (HI) assay using IDV strains from two viral lineages: D/bovine/Oklahoma/660/2013, D/660 lineage (hereby referred to as D/660); and/or D/swine/Italy/199724–3/2015 or D/bovine/France/5920/2014 (both of the D/OK lineage hereby referred to as D/OK).

French and Ukrainian samples were further tested as reported in table 1.

Samples from 3 dogs (3/122; 2.46% 95%CI 0.51-7.02%) and 1 cat (1/118; 0.85% 95%CI 0.02-4.63%) collected in Ukraine in 2024 tested positive for D/660, primarily from Odesa Oblast, except for one dog sample from Zaporizhzhia (table 2). All samples were negative for IAV by ELISA. All swab and tissue samples from France were IDV-negative by qRT-PCR.

**Table 2.**
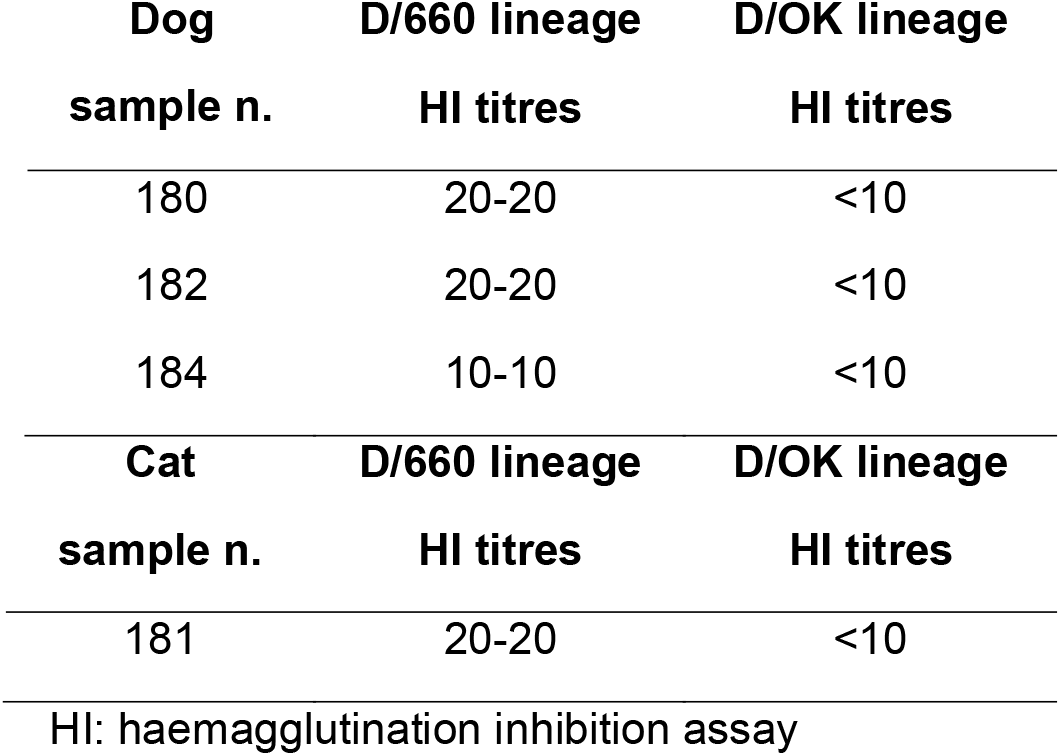
HI:Hae maggluti nation inhibitio n assay; IAV NP ELISA: influenz a A virus antibodi es by nucleopr otein ELISA.

Our findings provide evidence of IDV exposure in clinical healthy domestic cats (0.85%) and dogs (2.46%) from the Odesa and Zaporizhzhia Oblasts, Ukraine, although with relatively low rates and titres. These results align with two recent studies on IDV circulation in dogs and cats, reporting seroprevalence of 1.2% in 2016 and 4.7% in 2023 for D/660 in dogs in Southern Italy, and of 2.22% in cats in northern China (3, 4). In this study, household cats showed significantly higher exposure rates than stray cats, likely due to increased human contact, while the independent lifestyles of stray cats may limit exposure. Similar to our findings, the source of IDV infection in cats remains unclear. Although serology cannot confirm active transmission or clinical impact, the seroprevalence in domestic cats and dogs suggests that close human–animal interactions may increase the exposure risks. All the positive samples from 2024 belonged to the D/660 lineage, first detected in Europe in 2018 (5). None of these samples reacted against the D/OK lineage, that was likely replaced by D/660 lineage in Europe since 2019 (5).

The samples from pets in Italy and France tested negative for IDV. Since French samples were only tested against D/OK, a possible explanation is that this lineage was no longer circulating in France or, at least, in the surveyed area. Factors such as urbanization and the limited presence of cattle and other susceptible species, in this case, could be involved.

The French samples were also tested for ICV, and 2.63% of dogs tested positive, with HI titres between 20 and 80, supporting previous evidence of dog susceptibility. An old study conducted in France in 1988–1989 found HI reactivity in dogs as high as 32%, with the titres ranging from 1:20 to 1:320 (6).

The present study has some limitations. We used a convenience collection of samples, and in Italy and France, the samples were collected from a single geographic area. Also, demographic information was not available. Finally, different assays and IDVs were used for screening at the various locations. For instance, for the IDV screening, the French samples were tested only for D/OK, while the other samples were tested for both D/OK and D/660. Regarding ICV screening, only the French samples were tested.

Overall, our findings suggest that household dogs and cats might be exposed to IDV and could serve as a potential source of human infection. Proactive surveillance in pets is critical to understand IDV changing epidemiology and to mitigate potential public health concerns. In a One Health perspective, cats and dogs are uniquely positioned to act as reservoirs for IV infections in both household and in rural environments.

## Author Contributions

Conceptualization: CMT; formal analysis: CMT; investigation: AF, AF, AE, JO; resources: CMT, FP, MC, SLP, GM, MFD, NM, DM, EM; data curation: CMT, JO, MFD, SLP; writing—original draft preparation: CMT; writing—review and editing, MFD, VM, CM, FP, GL; AF visualization: CMT, AF; supervision: CMT, SLP, GM, MFD; project administration: CMT, SLP, GM, MFD. All authors have read and agreed to the published version of the manuscript.

## Funding

This publication was supported by the European Virus Archive GLOBAL project that received funding from the European Union’s Horizon 2020 research and innovation program (grant agreement no. 871029).

The research was supported by a grant from the National Research Foundation of Ukraine (Grant n. #2021.01/0006) and by the French agency for Research through the ANR FLUD (ANR-15-CE35-0005).

VM was supported by the National Laboratory for Infectious Animal Diseases, Antimicrobial Resistance, Veterinary Public Health and Food Chain Safety, RRF-2.3.1-21-2022-00001.

## Acknowledgement

The strain D/bovine/Oklahoma/660/2013 was kindly provided by Professor Feng Li from the University of Kentucky, USA.

The strain D/swine/Italy/199724–3/2015 was obtained from the European Virus Archive.

## Data Availability Statement

The original contributions presented in the study are included in the article; further inquiries can be directed to the corresponding author.

## Conflicts of Interest

EM is founder and Chief Scientific Officer of VisMederi srl.

## Ethics approval statement

The research has been waived by the Ethical Committee CESA-DiMeV, University of Bari, Italy (Protocol 5040-II/13 of 17 October 2024).

The animal study was reviewed and approved by the Institutional Animal Care and Use Committee of the National Scientific Center Institute of Experimental and Clinical Veterinary Medicine, Ukraine (Protocol #1-23, 19.04.2023).

Ethical review and approval were waived for the French samples, as they were collected and analyzed for diagnostic purposes.

## About the Author

Dr. Trombetta is an associate professor of hygiene and public health at the University of Siena. Her primary research interests include zoonotic viruses, infectious disease, and vaccine-preventable diseases.

---

**Supplemental figure 1** Geographical representation of the main areas across Europe where samples were collected. Red circles indicate regions where IDV positive samples were detected.

